# Patterns and environmental drivers of amphibian alpha and beta diversity in the Huangshan Mountains, southern Anhui, China

**DOI:** 10.64898/2026.08.20.745979

**Authors:** Suyue Wang, Ziyi Wang, Yujia Sun, Zhirong He, Qingyan Sun, Jiedong Wei, Yingxin Li, Meiting Liu, Jiayi Shi, Chunna Zhang, Siyu Wu, Yufeng Bai, Ziruo Zhang, Na Zhao, Supen Wang

## Abstract

Understanding the spatial patterns of biodiversity and their environmental determinants is fundamental to ecology and conservation, particularly in mountainous transitional zones where species assemblages can shift rapidly. Amphibians, as highly sensitive organisms, are excellent indicators of environmental change, yet quantitative assessments of their diversity in the Huangshan Mountains—a biodiversity hotspot at the junction of the Palaearctic and Oriental realms in southern Anhui, China—remain limited. We conducted systematic line-transect surveys across 200 grids (5 km × 5 km) during the spring and autumn of 2023 and 2024, recording a total of 10,342 individuals belonging to 23 species, 20 genera, 9 families, and 2 orders. Three species (*Fejervarya multistriata*, *Microhyla fissipes*, and *Bufo gargarizans*) were identified as dominant, and two nationally protected species (*Hoplobatrachus chinensis* and *Andrias davidianus*) were detected. Inter-annual alpha diversity did not differ significantly, but pronounced seasonal variation was observed, with spring supporting higher Shannon-Wiener and Pielou evenness than autumn in both years. Using generalized additive models and model averaging, we found that annual precipitation and the normalized difference vegetation index (NDVI) were consistently the strongest positive predictors of Shannon-Wiener diversity, Simpson dominance, and species richness, while species richness also declined significantly with increasing human footprint. Total beta diversity (Sørensen dissimilarity) was very high (0.981) and overwhelmingly driven by species turnover (97.86%) rather than nestedness. Partial Mantel tests and distance-based redundancy analysis further revealed that environmental distances—particularly annual precipitation—significantly shaped overall beta diversity and its turnover component after accounting for geographic distance. These results highlight the predominant role of climatic and vegetation gradients in structuring amphibian assemblages in this subtropical mountainous region, providing baseline data to inform local conservation strategies.

## 1 Introduction

Understanding the spatial distribution of biodiversity and the mechanisms that generate and maintain it remains a central pursuit in ecology and conservation biology(Edwards & Cerullo 2024; Wang et al. 2024). Species are not randomly distributed across landscapes; rather, their assemblages are shaped by a complex interplay of climatic conditions, topographic heterogeneity, habitat productivity, and anthropogenic pressures(Lawlor et al. 2024). Among vertebrates, amphibians are particularly sensitive to environmental variation owing to their permeable skin, physiological constraints, limited dispersal capacity, and dependence on both aquatic and terrestrial habitats(Riddell et al. 2026). These traits render them excellent biological indicators and ideal subjects for testing hypotheses about the environmental drivers of diversity patterns(Whitton et al. 2012). Because amphibian populations are declining worldwide due to habitat loss, climate change, pollution, and disease, spatially explicit assessments of their diversity have taken on renewed urgency(Luedtke et al. 2023).

Diversity can be examined through two complementary lenses(Tuomisto 2010). Alpha diversity captures local community structure in terms of species richness, evenness, and dominance, revealing how assemblages respond to site-specific environmental conditions. Beta diversity, in contrast, quantifies the degree of compositional dissimilarity among communities across space or environmental gradients. Partitioning beta diversity into its turnover and nestedness components offers mechanistic insights: high turnover implies species replacement along gradients, often driven by environmental sorting or dispersal limitation, whereas strong nestedness suggests ordered species loss, frequently associated with extinction or habitat filtering(Baselga 2010). Disentangling these components is essential to understand whether conservation actions should aim to protect environmental heterogeneity or maintain key habitats that support more impoverished communities(Tscharntke et al. 2012).

Mountainous regions, with their steep climatic gradients and structural complexity, constitute natural laboratories for biodiversity research(Rahbek et al. 2019). The Huangshan Mountain Range in southern Anhui, China, exemplifies such a setting. Situated on the northern border of the mid-subtropical zone, this area stands at the transitional boundary between the Palaearctic and Oriental zoogeographical realms(Shi et al. 2025a). The interaction of a subtropical monsoon climate, rugged topography, and extensive subtropical evergreen broad-leaved forests creates a mosaic of microhabitats that is expected to support diverse amphibian assemblages(Shi et al. 2025b). Despite its biogeographic significance and recognised status as a biodiversity hotspot, quantitative, spatially replicated studies that simultaneously analyse amphibian alpha and beta diversity and their environmental correlates in this transitional landscape remain scarce.

Previous work in other systems has identified several key environmental determinants of amphibian diversity, including temperature, precipitation, vegetation productivity, proximity to water bodies, and intensity of human disturbance(Qian et al. 2007). However, the relative importance of these drivers can vary substantially among biogeographic regions and spatial scales, and collinearity among predictors often complicates statistical inference(Bini et al. 2009). In the Huangshan Mountains, fragmentary faunal inventories exist, but a comprehensive, grid-based survey that controls for spatial autocorrelation and applies robust multivariate modelling to assess both alpha and beta diversity has not been conducted(Hong et al. 2025). This gap limits our ability to predict how amphibian communities in this transitional zone might respond to ongoing environmental change and to prioritise areas for conservation.

Here, we leverage systematically designed field surveys carried out during the peak activity seasons of 2023 and 2024 across 200 sampling grids in the Huangshan Mountains. Our objectives were threefold. First, we characterised amphibian species composition and alpha diversity patterns, examining both inter-annual stability and seasonal variation. Second, we quantified beta diversity and partitioned it into turnover and nestedness components to infer the processes underlying community differentiation. Third, we evaluated the independent and combined effects of climatic, topographic, vegetation, and human-disturbance variables on both alpha and beta diversity, using techniques that account for collinearity and spatial structure. Based on the region’s strong precipitation and productivity gradients, we hypothesised that annual precipitation and the normalized difference vegetation index would be the primary positive drivers of alpha and beta diversity, that the human footprint index would exert a negative influence, and that species turnover would dominate beta diversity given the environmental heterogeneity characteristic of the study area. By providing a robust, spatially explicit baseline, this study aims to inform amphibian monitoring and conservation planning in the southern Anhui mountainous area.

## 2 Materials and Methods

### 2.1 Study area

The study was conducted in the Huangshan Mountain Range, southern Anhui, China (29°59′–30°32′N, 117°12′–118°21′E), a biodiversity hotspot situated at the transition between the Palaearctic and Oriental zoogeographical realms(Liang 1998; Wang et al. 2022). The region has a subtropical monsoon climate with a mean annual temperature of 15–16 °C and annual precipitation of 1400–2200 mm(Wang et al. 2021). The topography is dominated by mountains and hills, and the zonal vegetation is subtropical evergreen broad-leaved forest(Wang et al. 2021). The study area covers approximately 5,500 km² across the districts of Huangshan, Guichi, Qingyang and Shitai.

### 2.2 Field surveys

Amphibian surveys followed the grid-based sampling framework of the national technical guidelines for county-level amphibian and reptile diversity assessment. A total of 200 grids, each 5 km × 5 km, were established across the study area(Therres et al. 2015). One line transect >200 m long and 2–4 m wide was placed in each grid, with a minimum distance of 5 km between adjacent transects(Cushman 2006; Rödel & Ernst 2004) (Figure 1). Surveys were conducted in spring (April–May) and autumn (September–October) of 2023 and 2024, corresponding to peak amphibian activity periods in this region(Liu et al. 2022; Pan et al. 2014). All surveys were carried out between 20:00 and 24:00 to coincide with the nocturnal activity of most amphibian species(Liu et al. 2022; Xiong et al. 2025). Each transect was walked at a speed of 2–3 km/h by a three-person team (two observers and one recorder). For each encounter, we recorded the species, number of individuals, geographic coordinates and elevation. Species identification followed Fei et al. (2006, 2009) and Fei et al. (2012), with nomenclature updated according to Frost (2024). The national conservation status of recorded species was determined using the National Key Protected Wild Animals List (2023), and threat categories were assigned based on the Red List of China’s Biodiversity(Jiang et al. 2021).

**Figure 1.**
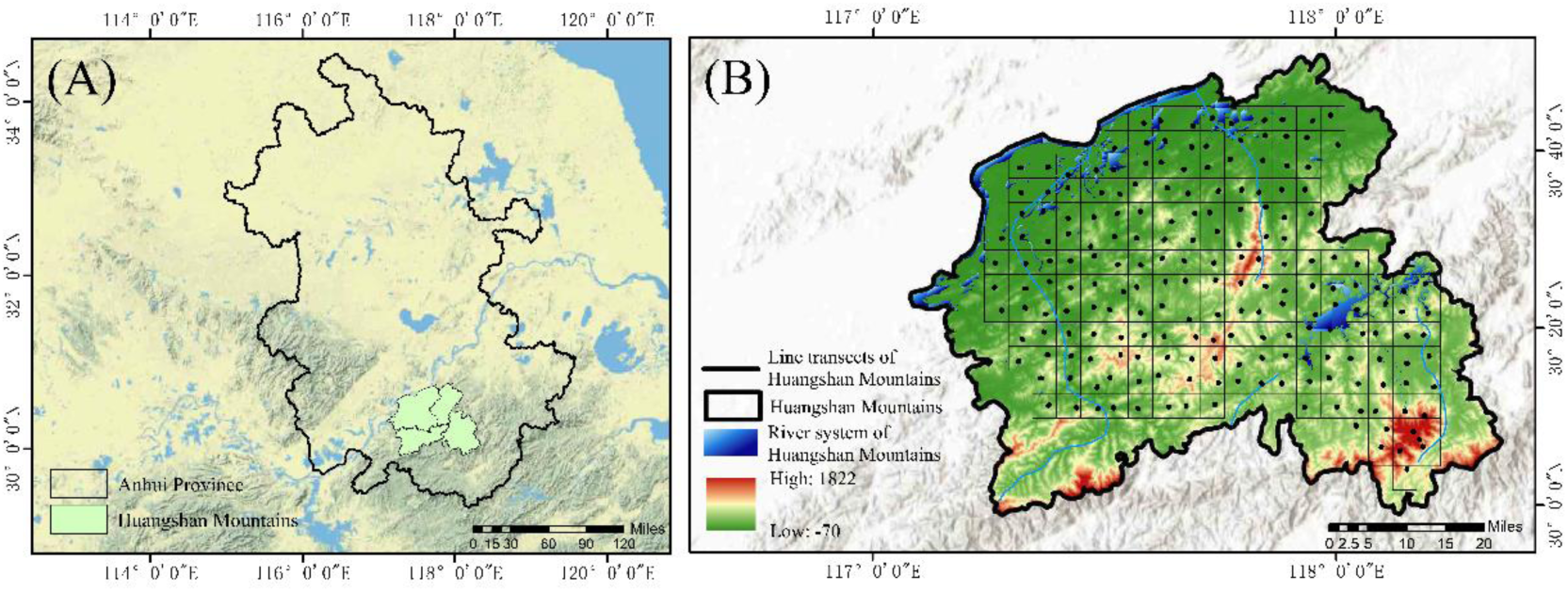
Location of the study area and distribution of amphibian survey transects in the Huangshan Mountains of southern Anhui Province. (A) Map of Anhui Province, highlighting the Huangshan Mountains in the southern part; (B) Topographic map of the Huangshan Mountains, indicating the 200 grids (5 km × 5 km) and the survey transects.

### 2.3 Environmental variables

We considered climatic, topographic, biotic and human-disturbance factors known to influence amphibian diversity(Duan et al. 2016; Ochoa-Ochoa et al. 2012; Zancolli et al. 2014). Two climatic variables (annual precipitation and mean annual temperature), three topographic variables (elevation, aspect and slope), two biotic variables (the Normalized Difference Vegetation Index, NDVI, and Euclidean distance to the nearest major river) and one anthropogenic variable (the Human Footprint Index) were obtained from public repositories (http://www.resdc.cn/) (Liu et al. 2022; Xiong et al. 2025). All layers were projected to the WGS 1984 UTM Zone 48N coordinate system and clipped to the Anhui provincial boundary in ArcGIS 10.8. Values of each variable were extracted for the midpoints of all 200 transects using the “Extract” tool.

### 2.4 Data analysis

All statistical analyses were performed in R version 4.5.2.

#### Alpha diversity and temporal variation

For each transect, species richness (C), Shannon-Wiener diversity index (H), Simpson dominance index (D) and Pielou evenness index (J) were computed. Dominant species were identified using the Berger-Parker dominance index (Pi = Ni/N), with thresholds of Pi ≥ 0.1 (dominant), 0.01 < Pi < 0.1 (common) and Pi ≤ 0.01 (rare)(Shi et al. 2025a). Differences in alpha diversity between years and between seasons within each year were tested with Mann– Whitney U tests(Gray et al. 2004).

#### Environmental predictors

To avoid collinearity, we calculated pairwise Spearman’s rank correlations among all 10 initially extracted variables. When |ρ| > 0.8, one variable of each pair was removed, giving priority to factors with clearer biological interpretation for amphibians(Xiao et al. 2016). Six non-collinear variables were retained: annual precipitation, NDVI, slope, aspect, distance to river and the Human Footprint Index.

#### Alpha diversity–environment relationships

Spearman correlations were first computed between each alpha diversity metric and the six predictors. Variables showing significant correlations (p < 0.05) were then entered into generalized additive models (GAMs). Variance inflation factors (VIF) were checked, and variables were retained only if VIF < 5.0(Zuur et al. 2009). Model selection was based on the corrected Akaike information criterion (AICc); when multiple models had ΔAICc < 2, model averaging was applied to the candidate set to obtain unconditional parameter estimates and variable importance weights(Burnham et al. 2004).

#### Beta diversity and partitioning

Pairwise Sørensen dissimilarity (βsor) was calculated from species presence/absence data. Total beta diversity was partitioned into turnover (βsim) and nestedness (βnes) components using the beta.pair function in the R package betapart(Baselga & biogeography 2010). The relative contribution of each component to total dissimilarity was expressed as a percentage.

#### Drivers of beta diversity

Geographic distances between transect midpoints were calculated with the distm function in the geosphere package, and environmental distances were computed as Euclidean distances on scaled environmental variables using vegdist in vegan. Partial Mantel tests (9999 permutations) were used to assess the relationship between βsor, βsim and βnes and each environmental distance matrix while controlling for geographic distance. Distance-based redundancy analysis (db-RDA) was additionally performed to quantify the unique and shared contributions of the six environmental predictors to beta diversity patterns, using the capscale function in vegan.

## 3 Results

### 3.1 Survey summary and species composition

Across the 200 transects, 10,342 amphibian individuals were recorded, belonging to 23 species, 20 genera, 9 families and 2 orders. Ranidae was the most speciose family (7 genera, 9 species, 39.13% of total richness), followed by Dicroglossidae (4 genera, 4 species, 17.39%). Microhylidae, Rhacophoridae and Salamandridae each contributed two species, whereas Bufonidae, Megophryidae, Hylidae and Cryptobranchidae were represented by a single species (Figure 2). Three species were dominant in the assemblage: *Fejervarya multistriata* (Pi = 0.43), *Microhyla fissipes* (Pi = 0.18) and *Bufo gargarizans* (Pi = 0.14). Two nationally protected Class-II species, *Hoplobatrachus chinensis* and *Andrias davidianus*, were detected. Species accumulation curves approached an asymptote in both years, indicating that sampling effort was sufficient (Figure 3).

**Figure 2.**
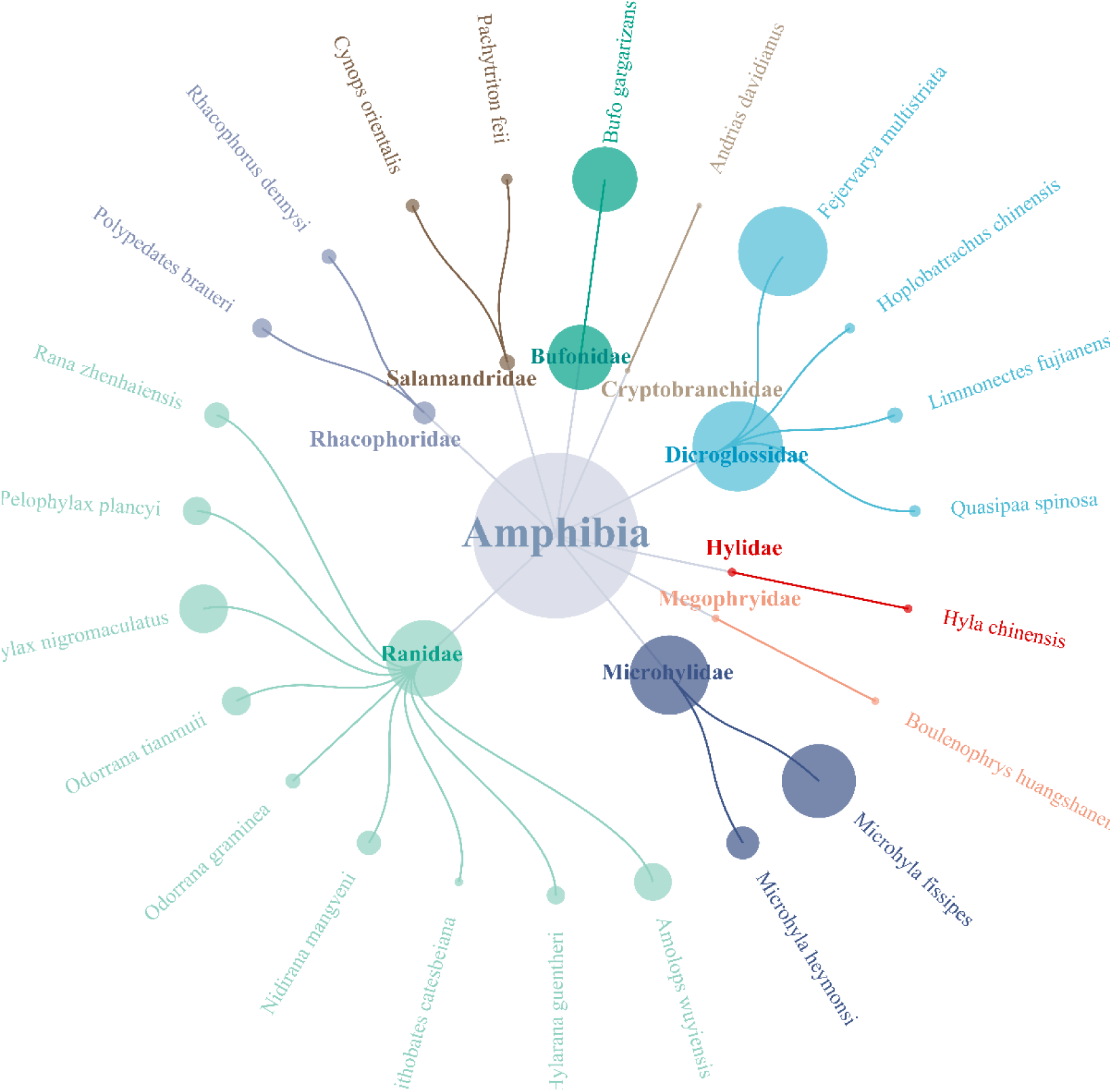
Species composition and relative abundance of amphibians in the Huangshan Mountains of southern Anhui Province. Circle size indicates relative abundance.

**Figure 3.**
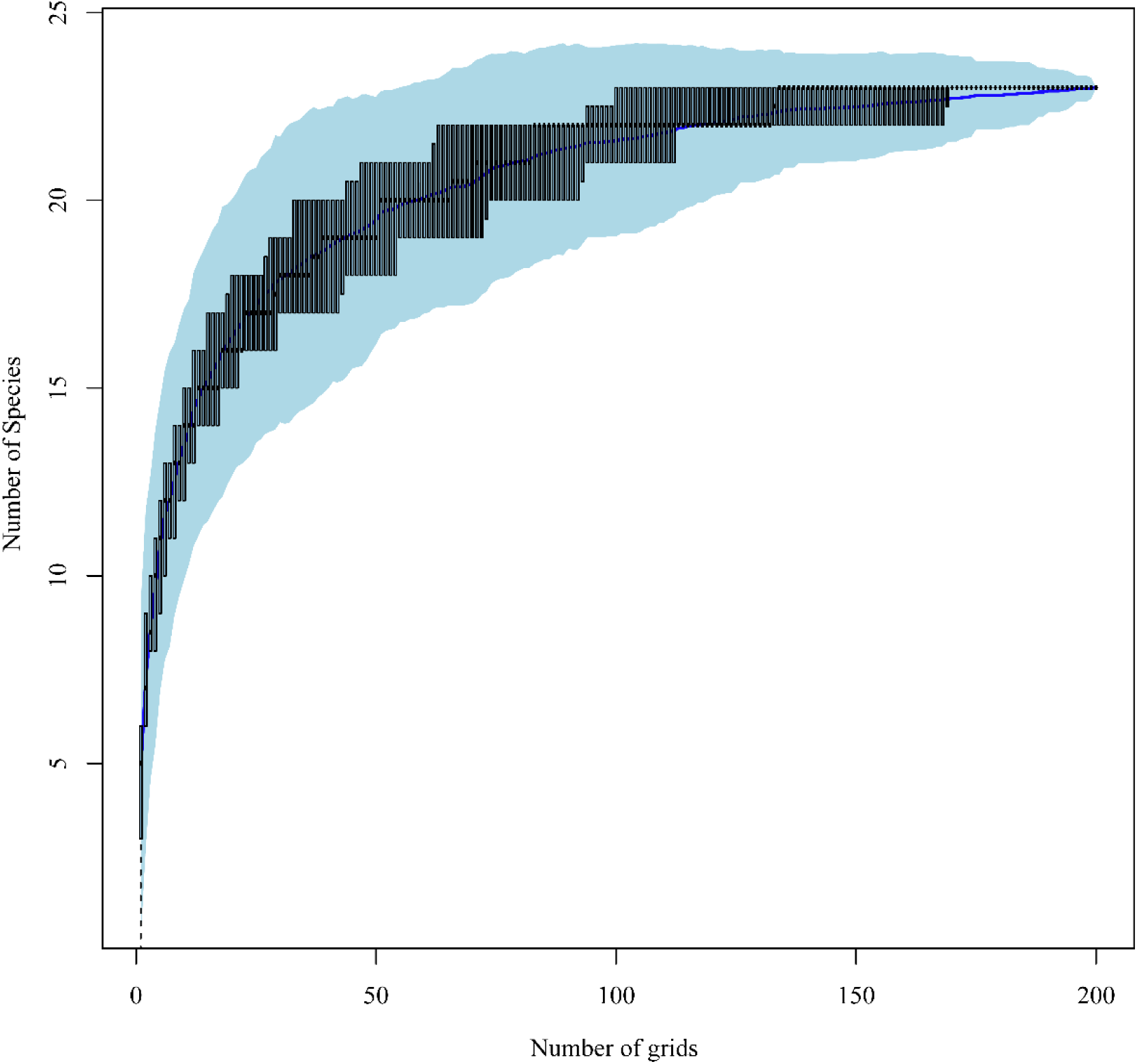
Species accumulation curves for amphibian surveys conducted in 2023 and 2024 in the Huangshan Mountains of southern Anhui Province.

### 3.2 Temporal variation in alpha diversity

Alpha diversity metrics did not differ significantly between 2023 and 2024 (Mann– Whitney U test, Shannon-Wiener: p = 0.55; Simpson: p = 0.66; Pielou: p = 0.36; species richness: p = 0.28). In contrast, pronounced seasonal variation was evident within each year. In 2023, Shannon-Wiener and Pielou evenness were significantly higher in spring than in autumn (p < 0.001), while Simpson index showed no seasonal difference (p = 0.26). In 2024, the same pattern held: spring values of Shannon-Wiener (p = 0.005) and Pielou evenness (p < 0.001) exceeded autumn values, and Simpson index again did not differ between seasons (p = 0.79) (Table 1).

**Table 1.**
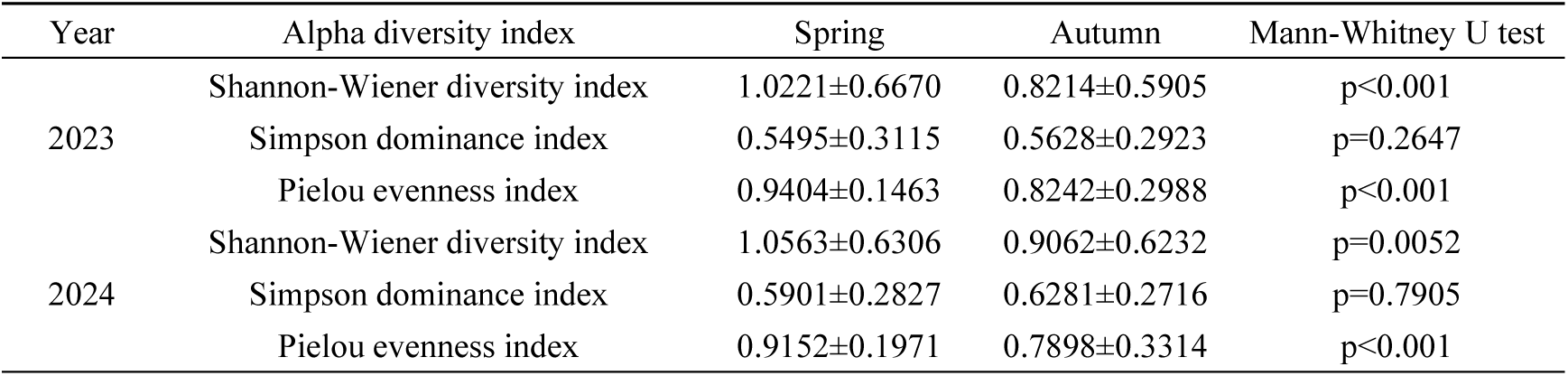
Inter-annual and seasonal differences in amphibian alpha diversity in the Huangshan Mountains of southern Anhui Province.

### 3.3 Alpha diversity–environment relationships

Strong collinearity was found among the original environmental variables; for instance, elevation was highly correlated with mean annual temperature and annual precipitation, and slope with terrain ruggedness. After removing correlated variables, the six retained predictors were annual precipitation, NDVI, slope, aspect, distance to river and the Human Footprint Index. Spearman correlations showed that Shannon-Wiener diversity, Simpson dominance and species richness were all significantly positively associated with annual precipitation, NDVI, slope and distance to river, and significantly negatively associated with the Human Footprint Index. No significant correlation was detected between any alpha diversity metric and aspect (Figure 4).

**Figure 4.**
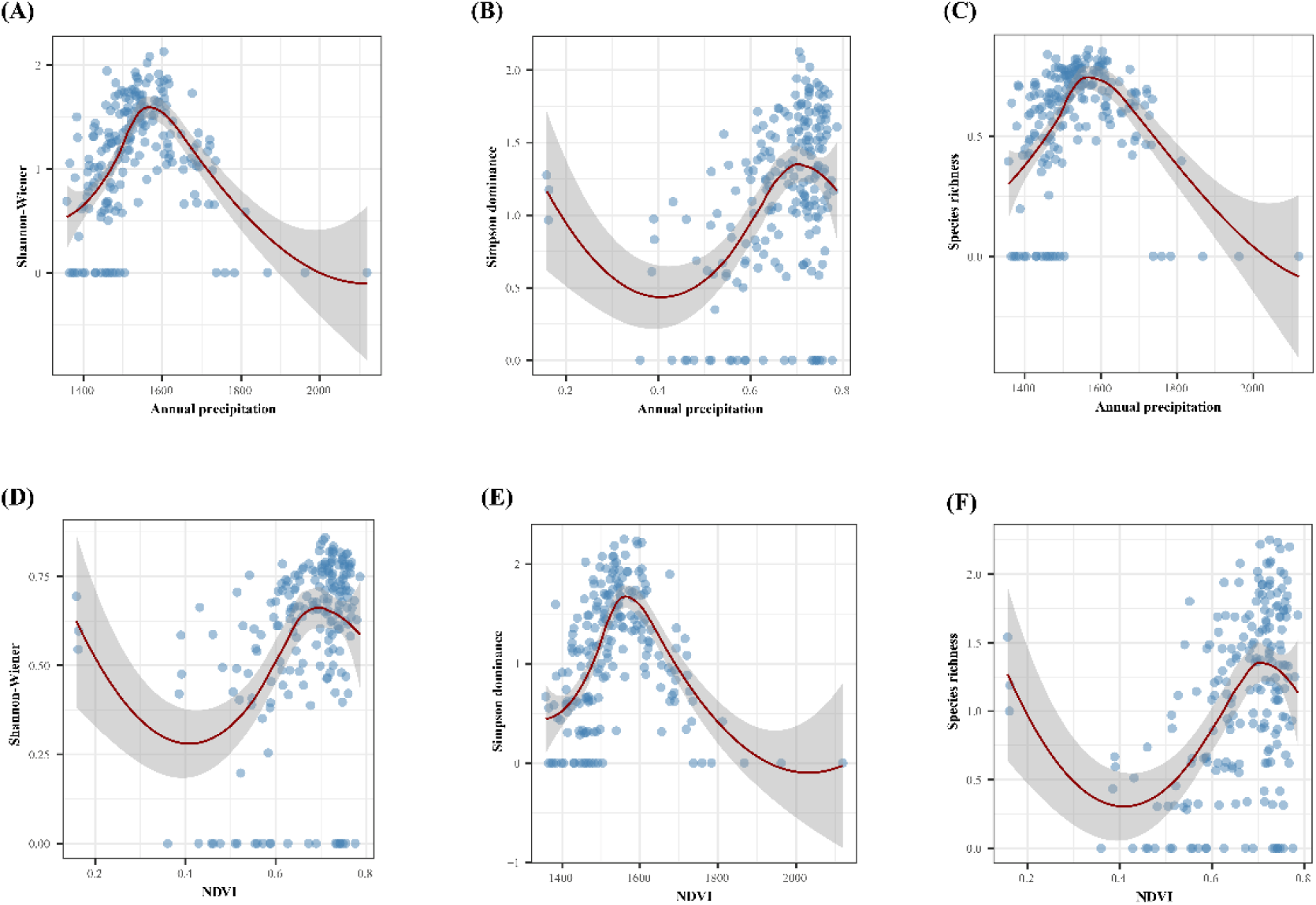
Effects of annual precipitation and NDVI on amphibian alpha diversity in the Huangshan Mountains of southern Anhui Province. (A), (B), and (C) show the relationships between annual precipitation and the Shannon-Wiener diversity index, Simpson dominance index, and species richness, respectively. (D), (E), and (F) show the relationships between the Normalized Difference Vegetation Index (NDVI) and the Shannon-Wiener diversity index, Simpson dominance index, and species richness, respectively. Blue dots represent observed values, red lines indicate the Spearman rank correlation fit, and shaded areas denote the 95% confidence intervals.

Generalized additive models and model averaging revealed that annual precipitation and NDVI were consistently the strongest predictors of alpha diversity (Table 2). Both variables were positively and significantly related to Shannon-Wiener diversity, Simpson dominance and species richness (all p < 0.001). In addition, species richness showed a significant negative relationship with the Human Footprint Index (p = 0.035), whereas this relationship was not significant for Shannon-Wiener or Simpson indices. The model weights for annual precipitation and NDVI were 1.0 (or close to 1.0) for all three alpha diversity metrics, confirming their predominant role. Other variables, including slope and distance to river, had lower and non-significant model-averaged coefficients (Table 2).

**Table 2.**
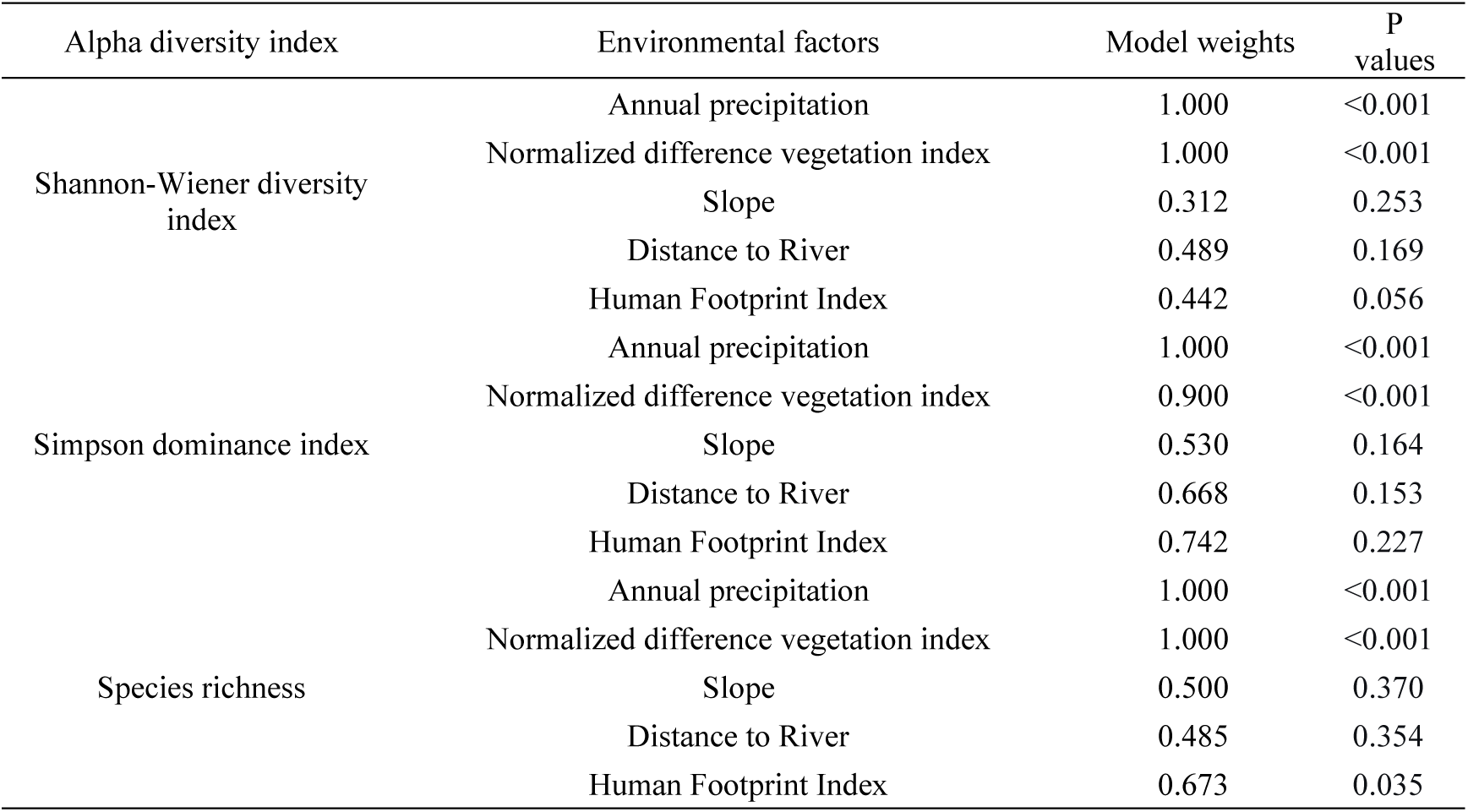
Model-averaged model weights and *P*-values for each alpha diversity index of amphibians in the Huangshan Mountains of southern Anhui Province.

### 3.4 Beta diversity patterns and drivers

Overall Sørensen dissimilarity among transects was 0.981, indicating strong compositional differentiation across the study area. Turnover (βsim = 0.960) accounted for 97.86% of total beta diversity, whereas nestedness (βnes = 0.021) contributed only 2.14%, demonstrating that community variation was overwhelmingly driven by species replacement rather than ordered species loss.

Partial Mantel tests, controlling for geographic distance, showed that total βsor was significantly positively correlated with environmental distances based on annual precipitation, NDVI and the Human Footprint Index (p < 0.01). The turnover component was significantly positively related to annual precipitation, NDVI and aspect (p < 0.05), while the nestedness component was significantly positively related to annual precipitation, slope and the Human Footprint Index (p < 0.05) (Figure 5).

**Figure 5.**
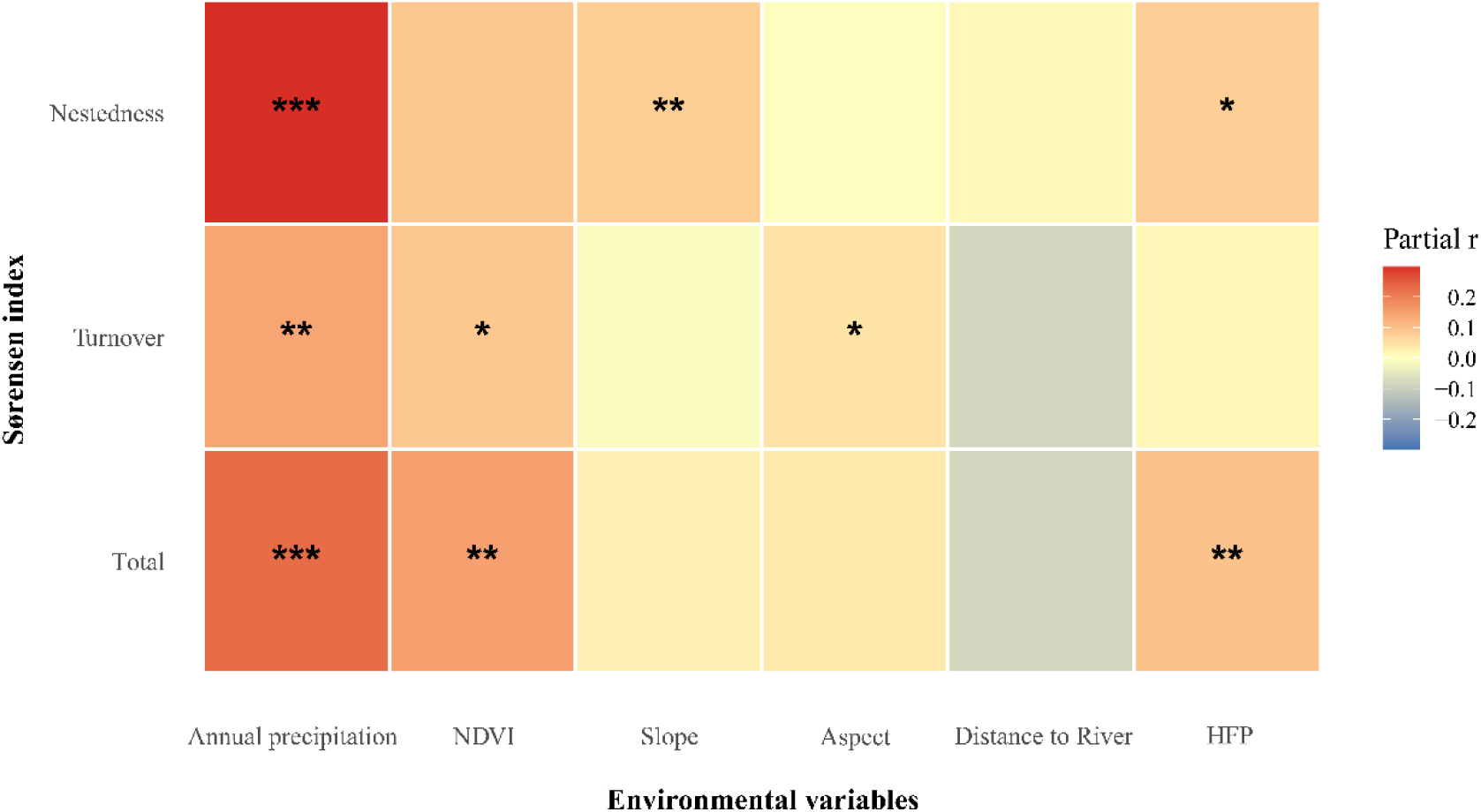
Relationships between amphibian beta diversity and environmental factors in the Huangshan Mountains of southern Anhui Province (*p < 0.05, **p < 0.01, ***p < 0.001).

Distance-based redundancy analysis further highlighted annual precipitation as the most influential predictor, with substantially higher explanatory power for both total βsor and the turnover component compared with the other environmental variables. Overall, these results identify annual precipitation as the key environmental gradient structuring amphibian beta diversity in the Huangshan Mountains, with species turnover being the dominant process.

## 4 Discussion

This study provides a spatially comprehensive assessment of amphibian diversity across the Huangshan Mountains, a biodiversity hotspot at the Palaearctic–Oriental transitional zone. Three main findings emerge: (i) amphibian alpha diversity was temporally stable between years but showed consistent seasonal variation, being higher in spring; (ii) annual precipitation and the normalized difference vegetation index (NDVI) were the primary environmental correlates of both alpha and beta diversity, while human footprint had a moderate negative effect on species richness; and (iii) amphibian community dissimilarity was exceptionally high and overwhelmingly driven by species turnover, not nestedness. These patterns align with the view that in topographically complex and climatically heterogeneous regions, species replacement along environmental gradients, rather than ordered species loss, structures communities. The observed seasonal shift in alpha diversity is ecologically intuitive (Tonkin et al. 2017). Spring surveys coincide with the peak breeding period for many amphibian species in subtropical China, when adults congregate at breeding sites and are more detectable; autumn surveys capture the post-breeding dispersal phase, often with juveniles dominating counts (Fe et al. 2010). The higher Shannon-Wiener and Pielou evenness values in spring therefore likely reflect both higher activity and a more equitable distribution of individuals among species during reproduction, whereas in autumn some species (e.g., *F. multistriata*) may become numerically overwhelming, lowering evenness without necessarily altering overall dominance (Simpson index). The lack of inter-annual difference indicates that, at least over the two-year period, the amphibian assemblages were stable, supporting the reliability of the data for spatial analysis, though longer-term monitoring would be required to detect directional changes.

Among the environmental factors examined, annual precipitation emerged as the most robust predictor of both alpha and beta diversity. This finding is consistent with numerous studies that identify water availability as a primary constraint on amphibian distributions (Buckley & Jetz 2007; Cunningham et al. 2016; Liao et al. 2025; Winter et al. 2016; Zhao et al. 2022). In the Huangshan Mountains, precipitation likely influences breeding-site hydroperiod, ambient humidity and the moisture of terrestrial microhabitats, all of which are critical for amphibian survival and reproduction. The positive association between NDVI and all alpha diversity metrics, and its significant link to beta diversity turnover, can be interpreted as the effect of vegetation productivity and structural complexity: dense, evergreen broad-leaved forests provide cooler, wetter microclimates and abundant invertebrate prey, supporting richer and more stable amphibian assemblages. Importantly, both precipitation and NDVI remained highly significant in multivariate models after accounting for other variables, and model-averaging gave them full or near-full model weights, underscoring their independent influence.

The Human Footprint Index, which integrates population density, land use and infrastructure, was negatively correlated with species richness, a pattern that corroborates the detrimental effects of human disturbance on amphibian communities (Venter et al. 2016). However, the relationship was significant only for species richness and not for Shannon-Wiener or Simpson indices. This may indicate that moderate disturbance can simplify communities by filtering out sensitive species without drastically altering overall diversity if common, disturbance-tolerant species (such as *F. multistriata*) remain abundant. The lack of a significant effect on evenness and dominance suggests that human footprint mainly acts by eliminating rare or specialized species, reducing richness, while the remaining dominant species maintain overall abundance patterns (Cazalis 2022; Hillebrand et al. 2008). No significant effect of slope, aspect, or distance to river was found in the ultimate models, implying that in this landscape, climatic and vegetation gradients may override local topographic and hydrographic factors at the grain of our analysis. However, these variables did show significant simple correlations with alpha diversity, so their influence is likely expressed through collinearity with precipitation and NDVI.

The beta diversity analysis revealed a striking predominance of the turnover component, accounting for nearly 98% of total dissimilarity. This result strongly suggests that amphibian communities are structured by environmental sorting across the strong precipitation and productivity gradients of the study area, rather than by dispersal limitation leading to nested species loss (Cunningham et al. 2016; Riddell et al. 2026). Such a pattern is expected in mountainous regions with high environmental heterogeneity (Hong et al. 2025). Partial Mantel tests controlling for geographic distance confirmed that environmental distances in annual precipitation, NDVI and human footprint independently explained total beta diversity, with precipitation again being the strongest correlate of turnover. The fact that turnover, not nestedness, drove the pattern implies that conservation efforts aimed at maintaining the full regional species pool should prioritize protecting the full range of environmental conditions— particularly the wetter, more-vegetated high-precipitation zones—rather than focusing only on a few species-rich sites (Wang et al. 2024).

Several limitations should be acknowledged. First, although the two years of surveys captured overall species richness well, as indicated by the accumulation curves, the temporal scope is insufficient to assess trends under climate change. Second, surveys were conducted exclusively at night, which may have reduced detection of species with diurnal or crepuscular activity in the surveyed season. Third, environmental variables were extracted at a 5-km grid resolution, which may mask fine-scale habitat features important for amphibians, such as the availability of temporary ponds or coarse woody debris. Future work could incorporate such microhabitat data and potentially use acoustic monitoring to enhance detection. Nevertheless, the strong and ecologically coherent relationships detected here provide a solid baseline.

In conclusion, this study demonstrates that amphibian diversity in the Huangshan Mountains is primarily structured by climatic moisture and vegetation productivity, with human disturbance playing a secondary role mainly via species loss. The dominance of species turnover in beta diversity highlights the conservation value of environmental gradients in this transitional mountain ecosystem. Protecting the remaining high-precipitation, forested areas from further human encroachment would be a priority action for safeguarding the region’s amphibian assemblages.

## Author Contributions

Suyue Wang: formal analysis (equal), investigation (equal), writing – original draft (equal). Ziyi Wang: investigation (equal). Yujia Sun: investigation (equal). Zhirong He: investigation (equal). Qingyan Sun: investigation (equal). Jiedong Wei: investigation (equal). Yingxin Li: investigation (equal). Meiting Liu: investigation (equal). Jiayi Shi: investigation (equal). Chunna Zhang: investigation (equal). Siyu Wu: investigation (equal). Yufeng Bai: investigation (equal). Ziruo Zhang: investigation (equal). Na Zhao: investigation (equal). Supen Wang: conceptualization (equal), formal analysis (equal), writing – original draft (equal).

## Acknowledgments

We thank the Amphibian and Reptile Survey Project in the Jianghuai Hilly Region and Huaihuai Plain of Anhui Province (2025BFAFZ02069-2)

## Ethics Statement

All animal experimental procedures received ethical review and approval from Anhui Normal University.

## Data Availability Statement

All datasets supporting the current study’s findings are provided within Data S1 in the Supporting Information.

