## Supplementary material for "Patterns and environmental drivers of amphibian alpha and beta diversity in the Huangshan Mountains, southern Anhui, China": table1

| Year | Alpha diversity index | Spring | Autumn | Mann-Whitney U test |
| --- | --- | --- | --- | --- |
| 2023 | Shannon-Wiener diversity index | 1.0221±0.6670 | 0.8214±0.5905 | p<0.001 |
|  | Simpson dominance index | 0.5495±0.3115 | 0.5628±0.2923 | p=0.2647 |
|  | Pielou evenness index | 0.9404±0.1463 | 0.8242±0.2988 | p<0.001 |
| 2024 | Shannon-Wiener diversity index | 1.0563±0.6306 | 0.9062±0.6232 | p=0.0052 |
|  | Simpson dominance index | 0.5901±0.2827 | 0.6281±0.2716 | p=0.7905 |
|  | Pielou evenness index | 0.9152±0.1971 | 0.7898±0.3314 | p<0.001 |
