## Supplementary material for "Patterns and environmental drivers of amphibian alpha and beta diversity in the Huangshan Mountains, southern Anhui, China": table2

| Alpha diversity index | Environmental factors | Model weights | | P values |
| --- | --- | --- | --- | --- |
| Shannon-Wiener diversity index | Annual precipitation | 1.000 | <0.001 | |
|  | Normalized difference vegetation index | 1.000 | <0.001 | |
|  | Slope | 0.312 | 0.253 | |
|  | Distance to River | 0.489 | 0.169 | |
|  | Human Footprint Index | 0.442 | 0.056 | |
| Simpson dominance index | Annual precipitation | 1.000 | <0.001 | |
|  | Normalized difference vegetation index | 0.900 | <0.001 | |
|  | Slope | 0.530 | 0.164 | |
|  | Distance to River | 0.668 | 0.153 | |
|  | Human Footprint Index | 0.742 | 0.227 | |
| Species richness | Annual precipitation | 1.000 | <0.001 | |
|  | Normalized difference vegetation index | 1.000 | <0.001 | |
|  | Slope | 0.500 | 0.370 | |
|  | Distance to River | 0.485 | 0.354 | |
|  | Human Footprint Index | 0.673 | 0.035 | |
